# Network and hierarchical organization of intrinsic timescales in the human brain

**DOI:** 10.64898/2026.07.01.735904

**Authors:** Bryan M. Krause, Emma F. Bublitz, Emily R. Dappen, Hiroto Kawasaki, Kirill V. Nourski, Matthew I. Banks

## Abstract

Intrinsic neural timescales represent the characteristic duration over which information is maintained in neuronal circuits. Evidence suggests that neural timescales vary systematically across the cortical hierarchy, with shorter timescales in primary sensory areas and longer timescales in higher-order association regions. In previous studies, hierarchy has been defined categorically, anatomically, or from the principal gradient of resting-state fMRI functional connectivity derived using diffusion map embedding (DME). Here, we assign hierarchical position to individual human intracranial electroencephalography (iEEG) recording sites by projecting their MNI coordinates onto this embedding space, derived from Human Connectome Project resting-state fMRI data. We estimated neural timescales from resting-state iEEG recordings in adult neurosurgical patients (n=46, 25 female) by extracting the aperiodic component of the local field potential power spectrum using spectral parameterization. Timescales increased monotonically with hierarchical position and associated with two region of interest (ROI)-level measures of network topology derived from DME of participants’ iEEG functional connectivity: ROIs with stronger mean functional connectivity exhibited longer timescales, as did ROIs functioning as hubs, defined by proximity to the center of embedding space. Finally, timescales varied with sleep stage, with slowest values during NREM and fastest during wake and REM. The hierarchical gradient present during wake and N1 was no longer detected during REM, N2, and N3 sleep, driven by a selective increase in timescales at lower levels of the hierarchy. This work presents a novel metric of hierarchy that can be applied to iEEG data, establishes a direct link between neural timescales, cortical hierarchy, and network topology in human iEEG, and demonstrates that this hierarchical organization is dynamically modulated by brain state.

**Significance Statement:** The brain processes information across multiple timescales simultaneously, with different cortical regions specialized for fast sensory processing or slow integration of context. We measured intrinsic neural timescales from human intracranial electrophysiological recordings and applied a novel method to locate each recording site along the cortical hierarchy. Timescales increased systematically from sensory to association areas, and were longest in network hubs, i.e., regions that are uniformly and widely connected to the rest of the brain. During sleep, this hierarchical organization was not detected, with sensory areas showing the largest increases in timescale. These findings advance our understanding of how the brain’s temporal organization is shaped by network architecture and modulated by brain state.

## Introduction

Neurophysiological signals comprise periodic (i.e., oscillatory) and aperiodic components. Oscillatory components of neuronal signals are postulated to mediate information transfer up and down the cortical hierarchy, with distinct frequency bands associated with direction-specific message passing (Bastos et al., 2012; Bastos et al., 2015), for example related to stimulus properties and prior expectation (Kumar et al., 2011; Sedley et al., 2016; Nourski et al., 2018). Here, we focus on non-oscillatory components of these signals, i.e., aperiodic activity, reflecting the endogenous dynamics of neuronal activity shaping this information transfer (He et al., 2010).

Intrinsic neural timescales derived from resting state aperiodic activity have emerged as biomarkers of brain dynamics (Golesorkhi et al., 2021b; Wolff et al., 2022). Timescales are readily measurable under task-free (resting state) conditions from blood-oxygen-level-dependent (BOLD) fMRI, scalp EEG, local field potentials (LFPs), and single unit activity (Wolff et al., 2022; Zeraati et al., 2023)(SUA) using the monoexponential decay of the autocorrelation function, or, equivalently, the ‘knee frequency’ of the power spectral density (Chaudhuri et al., 2018; Gao et al., 2020). By describing the autocorrelation (self-similarity over time) of neural signals (Honey et al., 2012; Murray et al., 2014), neural timescales reflect the process of neuronal temporal integration (Gao et al., 2020), which is key to perceptual and cognitive function (Stephens et al., 2013; Norman-Haignere et al., 2022; Giroud et al., 2024). Neural timescales vary with aging, reflecting both cellular and circuit-level alterations (Voytek et al., 2015; Gao et al., 2020; Wu and Gollo, 2025), and neurological and psychiatric conditions such as autism, aphasia, dementia, and psychosis (Watanabe et al., 2019; Wengler et al., 2020; Uscătescu et al., 2023; Murai et al., 2024; Zhang et al., 2024; Djimbouon et al., 2025). Furthermore, evidence suggests that timescales adapt dynamically to cognitive and perceptual demands (Gao et al., 2020; Zeraati et al., 2023; Trepka et al., 2024; Raposo et al., 2025; Çatal et al., 2025). For example, timescales are dynamically modulated during visual attention and working memory (Zeraati et al., 2023; Raposo et al., 2025).

The brain is organized hierarchically (Felleman and Van Essen, 1991; Douglas and Martin, 2004; Murray et al., 2014), and several previous studies have shown that timescales vary systematically across the cortical hierarchy, with short timescales in sensory cortex and longer timescales in association cortex (Murray et al., 2014; Gao et al., 2020; Golesorkhi et al., 2021b; Wolff et al., 2022). Shorter neural timescales could mediate encoding of rapidly changing stimulus features processed by early sensory areas, whereas longer neural timescales could enable sustained integration and contextual inference in association areas (Honey et al., 2012). For example, the time window over which neurons integrate sensory inputs lengthens along the auditory cortical hierarchy (Overath et al., 2008; Norman-Haignere et al., 2022; Norman-Haignere et al., 2025), and timescales vary by region with auditory response latency (Cusinato et al., 2023). This hierarchical process of “zooming-out” in time, allowing integration over progressively broader time scales, parallels the expansion of visual receptive fields in higher visual cortex(Shafiei et al., 2020).

In practice, measuring the cortical hierarchy quantitatively is challenging, especially in the human brain. Data from anatomical tracer studies provide the clearest quantitative description of hierarchy in preclinical models (e.g., (Markov et al., 2014)) but are unavailable in human subjects. The ratio of T1w/T2w magnetic resonance images has been used as a surrogate (Burt et al., 2018; Gao et al., 2020; Ito et al., 2020; Cusinato et al., 2025), as it reflects the density of myelin, which is expected to be higher in sensory compared to association cortex (Glasser and Van Essen, 2011; Glasser et al., 2014), but this measure is indirect. Another promising approach is one based on the *functional* hierarchical organization of cortical and subcortical networks, where hierarchy emerges as a gradient of similarity of regional connectivity to the larger brain (Coifman et al., 2005; Margulies et al., 2016; Shafiei et al., 2020). Here, we extend this approach to intracranial electrophysiological recordings, as we have done previously (Banks et al., 2023), and estimate the hierarchical position of individual recording sites. Resting state connectivity from the Human Connectome Project was mapped into an embedding space in which distance represents functional dissimilarity in connectivity to the rest of the brain. This procedure yields a hierarchical gradient of brain regions. Each brain region has a corresponding spatial coordinate in the MNI template brain. We then use the MNI coordinates of recording sites to map them onto this hierarchy.

## Methods

### Ethics statement

Research protocols were approved by the University of Iowa Institutional Review Board. Written informed consent was obtained from all participants. Participation in research did not interfere with clinically necessary care, and participants were free to withdraw consent at any time without consequence to their clinical management.

### Participants

Participants were adult neurosurgical patients (*n*=46; 25 female) at the University of Iowa Hospital and Clinics (UIHC) with medically refractory epilepsy undergoing chronic iEEG monitoring to localize seizure foci prior to resection surgery. Data collection took place within the years 2017 to 2026. Each participant underwent audiometric and neuropsychological testing prior to electrode implantation. Exclusion criteria included moderate hearing loss, severe cognitive impairment, and previous resection. Experimental sessions took place in electrically shielded suites in the Epilepsy Monitoring Unit. Participants were tapered off their antiepileptic drugs during the monitoring period when experimental data were collected.

### Electrophysiological recordings

Resting state (i.e., task-free) iEEG recordings were obtained while participants lay quietly in their hospital bed. Daytime recording sessions took place a median (interquartile range [IQR]) of 6.8 (3.9 – 8.0) days after electrode implant surgery. Recordings were obtained using depth multicontact stereo EEG arrays (sEEG, *n* = 2494 sites) and subdural grids (*n* = 2932 sites). All electrodes were placed solely based on clinical requirements, as determined by a team of epileptologists and neurosurgeons. iEEG data were collected using a Neuralynx Atlas System (Neuralynx, Bozeman, MT, USA), amplified, filtered (0.1-500 Hz bandpass, 5 dB/octave rolloff), and digitized at 2 kHz.

### Sleep recordings

Overnight sleep recordings were performed in 27 participants without clinical contraindications who consented to additional scalp electroencephalography and electromyography electrodes needed for polysomnography. Sleep recordings took place a median (IQR) 7.5 (6.3 – 8.3) days after electrode implantation surgery. A median (IQR) of 519 (429—596) min of overnight data was used per participant. Sleep stages (wake, N1, N2, N3, and REM) were scored manually in 30-s epochs according to standard American Academy of Sleep Medicine criteria (Berry et al., 2017).

### Anatomical reconstruction

Recording sites were localized as in our previous work (Banks et al., 2023) based on pre- and post-implantation T1-weighted structural MRI and post-implantation computed tomography (CT). Briefly, post-implantation images were aligned to pre-implantation MRI using linear coregistration as implemented in FSL (FLIRT) (Jenkinson et al., 2002). Electrodes were manually identified as magnetic susceptibility artifacts or as metallic hyperdensities in post-implantation MRI and CT, respectively. To correct for post-operative brain shift and distortion, electrode locations were further refined within the space of the pre-operative MRI using three-dimensional non-linear thin-plate spline warping (Rohr et al., 2001). Electrode locations were then mapped into a common anatomical template space (MNI-152), involving linear coregistration of the T1-weighted MRI to the MNI-152 template (1-mm resolution) with FSL’s FLIRT and application of the resulting transformation to the electrode coordinates in participant space.

### iEEG preprocessing

Preprocessing of iEEG data was performed with custom functions written in MATLAB (MathWorks, Natick, MA, USA) as described previously (Krause et al., 2023). Recording sites in the cortical gray matter, hippocampus, and amygdala were used for analysis. After initial rejection of recording sites identified as seizure foci, automated preprocessing steps excluded recording channels or time intervals with atypical power in any band, transient voltage deflections, or amplifier clipping. Shared broadband noise across channels was removed by constructing a spatial filter from singular value decomposition of the high-frequency (>200 Hz) signal, then applying this filter to the broadband signal to remove the common signal. Further information on this procedure can be found in our prior work (Banks et al., 2023; Krause et al., 2023). LFP data were downsampled to 1 kHz and de-noised using the demodulated band transform (DBT)(Kovach and Gander, 2016) and divided into 60-sec segments for analysis.

### Calculation of neural timescales

Power spectral densities (PSDs) were estimated from artifact-free 60-s resting-state segments using Welch’s method (MATLAB function “pwelch”, Hamming window with 8 segments overlapped by 50%, and with nfft = 2048). Timescales were estimated from Lorentzian function fits (spectral parameterization, aka ‘FOOOF’) to the PSD (Donoghue et al., 2020) over the range 0.1–100 Hz with aperiodic_mode = “knee”, peak_width_limits = [1,20], max_n_peaks = 5, min_peak_height = 0, and peak_threshold = 2.

The intrinsic timescale was computed from the fitted knee frequency f_K_ as:

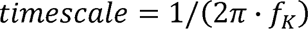

Timescales were log-transformed for analysis. Although the relationship between timescale and knee frequency is non-linear, the relationship between log timescale and log frequency is linear (with slope = −1). Therefore, all analyses of log timescale are linearly equivalent to analyses on a log frequency scale.

### Diffusion map embedding (DME) analysis

To test hypotheses about the hierarchical organization of neural timescales, we used DME (Coifman and Hirn, 2014; Margulies et al., 2016) as in our previous work (Banks et al., 2023; Krause et al., 2023).

Group average resting state connectivity from the WU-Minn Human Connectome Project S1200 data release (Van Essen et al., 2013) was parcellated with the Schaefer-Yeo 1000-region cortical atlas (Yeo et al., 2011; Schaefer et al., 2018) and the Tian S2 32-region subcortical atlas (Tian et al., 2020). Resulting parcels were mapped into an embedding space in which distance represents functional dissimilarity in connectivity to the rest of the brain. Specifically, functional connectivity matrices were thresholded using k-nearest neighbors to retain at minimum the largest 10% of connections in each row while ensuring the minimum spanning tree is included (an ‘OR’ operation). Cosine similarity was calculated across rows of the thresholded matrix to produce a similarity matrix **K** which was then normalized by degree matrix **D** to give a symmetric matrix **P_symm_**= **D**^−0.5^**KD**^−0.5^. The embedding space results from eigendecomposition of **P_symm_**. We used the multiscale implementation to consider simultaneously all values for the diffusion parameter *t* (Richards et al., 2009).

We defined a hierarchical gradient along a vector in embedding space pointing from the mean N-dimensional coordinate across visual and somatomotor ROIs, defined as the bottom of the hierarchy, toward the mean N-dimensional coordinate across default mode ROIs, defined as the top of the hierarchy. To assign a continuous representation of hierarchy, we projected the N-dimensional data along that gradient vector and applied min-max scaling to produce Hierarchy values strictly between 0 and 1 (see **Fig. 2**). To assign Hierarchy values to iEEG recording sites, we mapped spatial recording site coordinates within the MNI-152 template brain to the nearest cortical or subcortical voxel in the Schaefer-Yeo 1000-region or Tian S2 32-region atlas, respectively.

**Figure 1.**
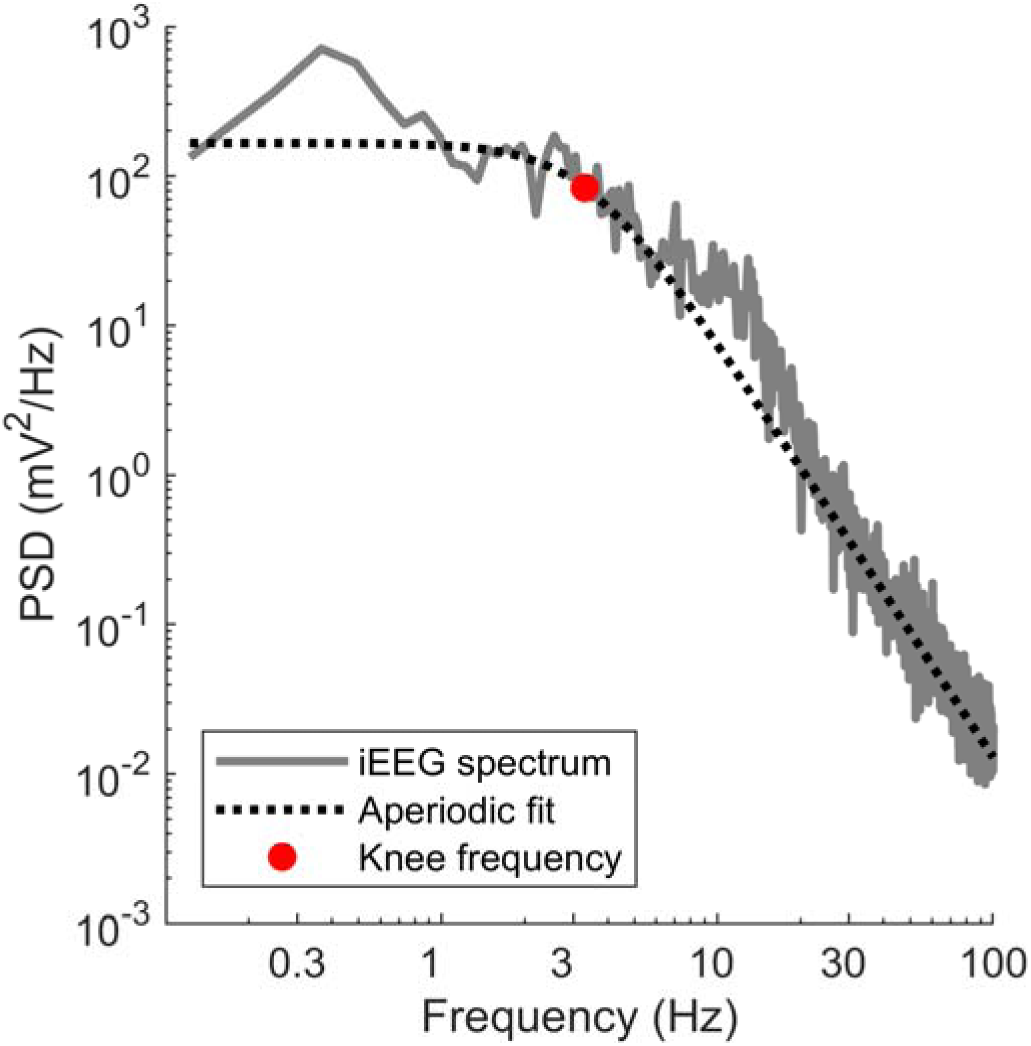
Measurement of the intrinsic timescale. Resting state power spectral density (PSD) from one recording site (orbitofrontal gyrus; grey line) and superimposed aperiodic fit (dashed line) with knee frequency indicated (red circle). The intrinsic timescale was computed as the reciprocal of (2π × knee frequency); in this example the knee frequency is 3.3 Hz and the intrinsic time constant is 48 ms.

**Figure 2.**
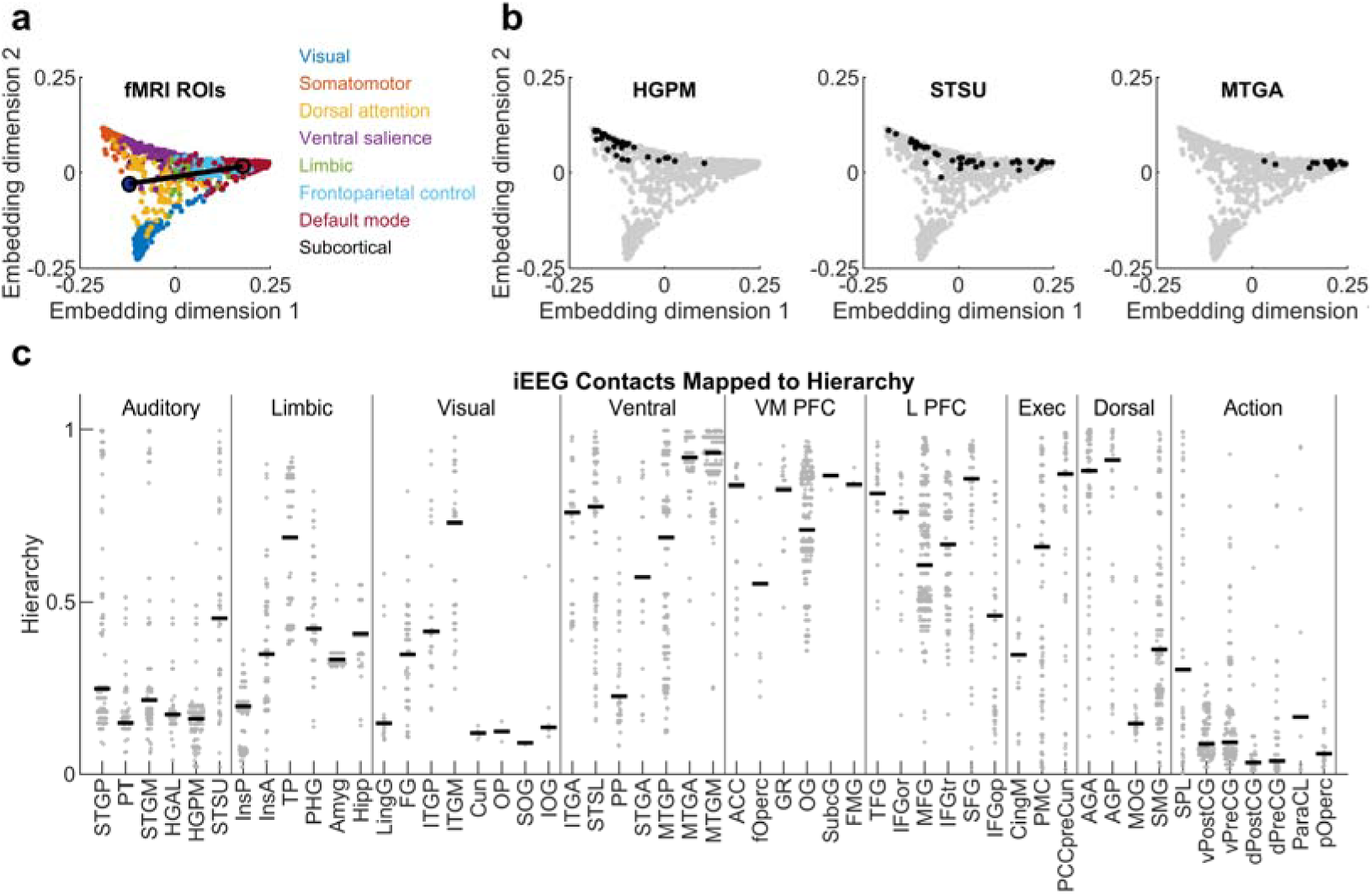
Measurement of the cortical hierarchy. **A.** fMRI BOLD resting state connectivity data from the human connectome project were mapped into an embedding space in which distance represents functional dissimilarity (in connectivity to the rest of the brain). Data from 1000 ROIs are shown plotted in the first two dimensions of embedding space and color-coded according to canonical resting state networks. A hierarchy is apparent from primary sensory areas (blue) to the default mode ROIs (red). **B.** iEEG recording sites in 48 participants were matched to fMRI-derived ROIs using MNI coordinates. Black dots show the corresponding locations in embedding space after mapping for iEEG contacts previously assigned to HGPM, STSU, and MTGA (left, center, right, respectively). **C.** Mapped hierarchy of iEEG contacts by ROI. Functional ROI groupings (Auditory, Limbic, etc.) were derived in previous work (Banks et al., 2023). Gray dots are individual recording sites, black horizontal lines indicate median across recording sites within each ROI.

A separate DME analysis was used to characterize network topology from the iEEG data (**Fig 4**). Functional connectivity was computed between recording sites within each participant as orthogonalized gamma band-power envelope correlations as in our previous work (Banks et al., 2023); orthogonalization discounts connectivity near zero phase lag and thereby limits volume-conduction artifact. Connectivity was estimated from 60-s segments of data, averaged within ROIs to form an ROI × ROI matrix per participant, and averaged across participants to yield a grand-average matrix on which DME was performed. From this embedding, we used the proximity of an ROI to the center of embedding space as a measure of “hubness” (smaller distance = more hub-like; (Banks et al., 2023)) alongside mean functional connectivity.

**Figure 3.**
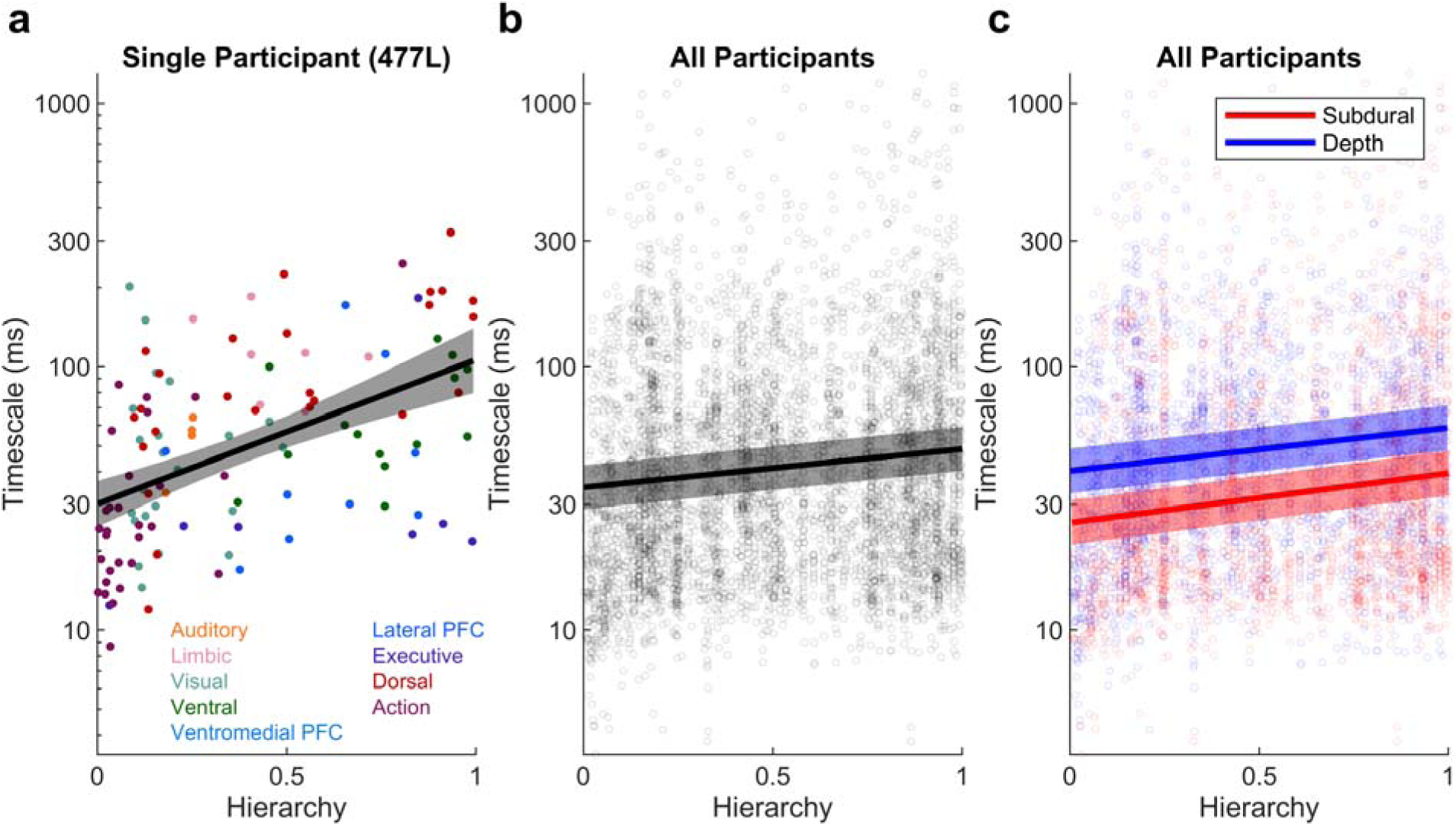
Hierarchical organization of neural timescales. **A.** Timescales measured from power spectral densities in a representative participant, labeled according to our previous categorization of iEEG ROIs into functional groups (Banks et al., 2023). **B.** Timescales measured from all 5426 recording sites (N=46 participants). **C.** Recording sites labeled by recording type: either subdural surface grids or depth (sEEG). For all panels, each point is a single recording site. Regression lines and 95% confidence intervals are from mixed effects models (or simple linear regression for panel A).

**Figure 4.**
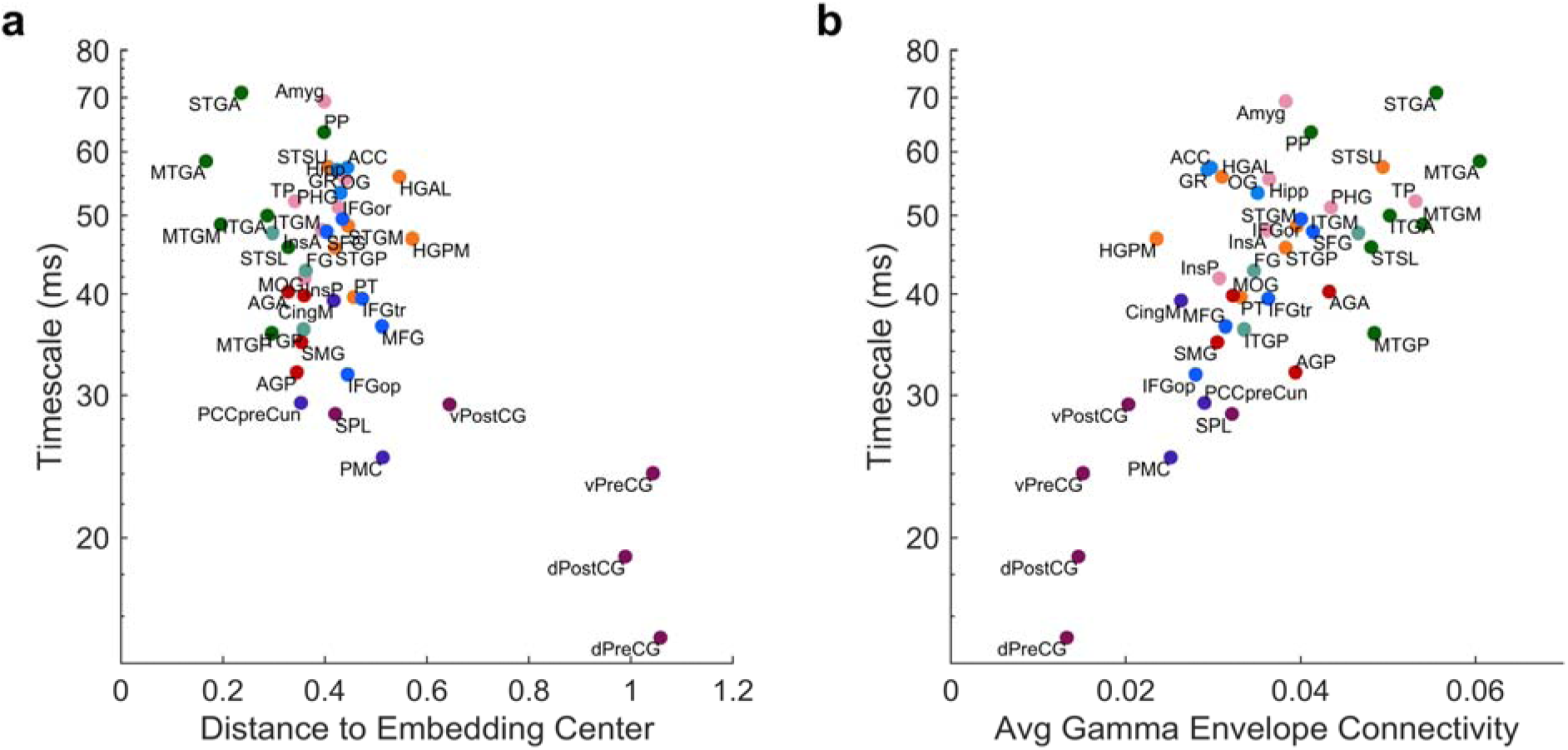
Neural timescales vary with network connectivity. A. Timescales measured from resting state power spectral density are negatively correlated with distance to the center of the DME (i.e., “hubness”; Ref.(Banks et al., 2023)), and B. positively correlated with mean gamma-band connectivity (right).

### Statistical modeling

We used linear mixed-effects models (LME) to model the effects of Hierarchy, Contact Type, Region-of-interest, and/or Sleep Stage (fixed effects) while accounting for the nested structure of the data within participants and within recording sites (random effects).

For RS analyses we modeled the median timescale across segments (resulting in a single observation per recording site), so the LME model was:

log_10_(Timescale) ∼ Hierarchy + (1|participant)

For presentation of single-participant data, we also present simple regression fits of log_10_(Timescale) ∼ Hierarchy.

We also present a model that adjusted for Electrode Type (two-level factor: depth or subdural) and allowed for different slopes and intercepts with Hierarchy per Electrode Type:

log_10_(Timescale) ∼ Hierarchy * ElectrodeType + (1|participant)

For comparisons with DME-derived estimates of ‘hubness’, we used a mixed effects model to produce an average log-timescale for each ROI, adjusted for sampling within participants:

log_10_(Timescale) ∼ ROI + (1|participant)

To model differences in the relationship between timescales and hierarchy in sleep, we fit a model allowing for different slopes and intercepts by sleep stage. For this model, we took the median timescale across segments per sleep stage, so there are multiple observations per recording site (one per stage) and we include a random intercept for recording site:

log_10_(Timescale) ∼ Hierarchy * SleepStage + (1|participant) + (1|recording site)

For all models, we interpreted the results by making model predictions at the marginal level (averaged over the random effects) with confidence intervals determined from the uncertainty in the fixed effects, representing an estimate of the population average relationship. Predictions on the original (ms) scale were adjusted for bias in the back-transformation from log scale. Because the response variable was modeled on log_10_ scale, we exponentiated contrasts with base 10 to report comparisons at a given Hierarchy value as fold change. For differences in slope with Hierarchy this transformation represents the marginal ratio of timescales high versus low on the hierarchy which we label hierarchical timescale ratio (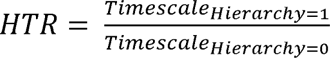).

## Results

### Measuring intrinsic neural timescales

Intrinsic neural timescales were estimated for 5426 recording sites (N = 46 participants; see Table 1) from resting-state iEEG power spectral densities (PSDs) using the spectral parameterization (formerly “FOOOF”) framework (Donoghue et al., 2020). Neural power spectra recorded under task-free conditions are well characterized by a combination of aperiodic (“1/f-like”) broadband activity and superimposed narrowband oscillatory peaks. Spectral parameterization decomposes these components by fitting a Lorentzian function to the aperiodic component, which takes the form of a spectral “knee” whose frequency reflects the characteristic timescale of the underlying neural dynamics. The intrinsic timescale was computed as Timescale = 1 / (2π × knee frequency). A representative example is shown in Figure 1, illustrating a PSD from a single recording site in orbitofrontal gyrus.

**Table 1.**
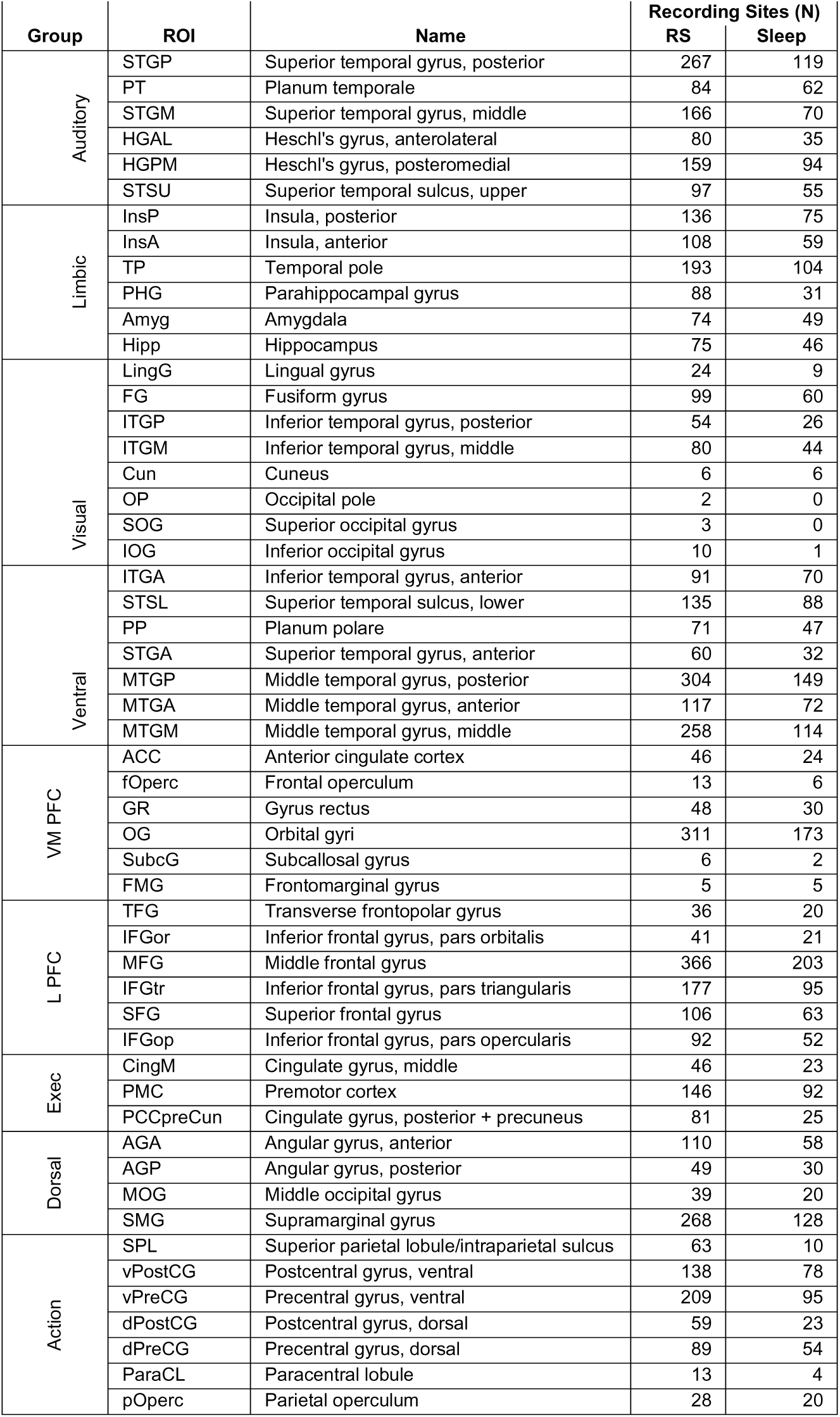
Recording site coverage. VMPFC = ventromedial prefrontal cortex. L PFC = lateral prefrontal cortex.

### Measuring functional hierarchy

To quantify the functional hierarchical position of each iEEG recording site, we employed diffusion map embedding (DME) of resting-state functional connectivity (Margulies et al., 2016; Banks et al., 2023; Krause et al., 2023). Group-average BOLD fMRI resting-state functional connectivity from the Human Connectome Project was projected into an embedding space in which Euclidean distance between regions reflects their dissimilarity in whole-brain connectivity. We defined a hierarchy vector directed from the centroid of visual/somatomotor to the centroid of default mode ROIs which lay mostly along the first embedding dimension/gradient (Figure 2a). Hierarchy position estimates were assigned to iEEG recording sites by finding the nearest fMRI-derived ROI in MNI space and projecting along the hierarchy vector. The mapped hierarchy of iEEG contacts recapitulates the expected functional organization, with a progression from low hierarchical position in primary sensory and auditory regions (e.g., Heschl’s gyrus, primary motor cortex) to high hierarchical positions in transmodal areas (e.g., middle temporal gyrus, prefrontal cortex, and cingulate cortex) while also allowing hierarchy to vary with position within those previously assigned ROIs (Figure 2b,c). We use this DME-based hierarchy index to measure the relationship between hierarchy and neural timescales in subsequent analyses.

### Resting-state relationship between neural timescales and hierarchy

Neural timescales increased with hierarchical position in resting-state iEEG data recorded during daytime wakefulness in an example individual participant (Figure 3a) and across all participants (Figure 3b). We characterized the log-scale slope between hierarchy and timescale using model marginal predictions of the ratio between timescales high and low on the hierarchy (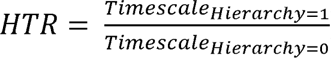). For resting state, the HTR was 1.40, 95% CI [1.33, 1.47]. Estimates of timescale were longer for depth versus subdural recording sites at a given level of the hierarchy (Figure 3c; at Hierarchy = 0.5: subdural 32 ms, 95% CI [26, 39]; depth 49 ms, 95% CI [40, 59]), but the slope with hierarchy was not significantly different (coefficient for Subdural*Hierarchy = 0.024, 95% CI [-0.020, 0.069], p = 0.29).

We previously used DME to map the functional organization of resting-state networks recorded intracranially (Banks et al., 2023), and sought to relate that functional organization to measured neural timescales. Replicating our previous method, resting-state iEEG functional connectivity (orthogonalized gamma-band envelope correlation) was computed at the single-participant level, averaged within ROIs to yield ROI × ROI matrices per participant, and then averaged across participants to produce a grand-average connectivity matrix on which DME was performed. We then characterized ROI “hubness” in two dimensions: proximity to the center of the embedding space and mean functional connectivity strength (Banks et al., 2023). A region close to the embedding center exhibits a connectivity profile maximally similar to the whole-brain average, marking it as a functional network hub. ROI-level timescales showed a strong negative correlation (Pearson correlation = -0.58, p < 0.001) with distance from the embedding center (Figure 4a): hub-like ROIs exhibited the longest timescales, while ROIs at greater distances showed faster dynamics. In parallel, timescales were strongly and positively correlated (Pearson correlation = 0.65, p < 0.001) with mean functional connectivity strength (Fig. 4b): ROIs with the strongest average coupling to the rest of the network exhibited the slowest dynamics, while those with the weakest mean connectivity showed the fastest timescales. Together, these associations indicate that timescales are closely linked to the functional network architecture of the cortex, with slower timescales in regions that are both more central within the network geometry and more strongly coupled with the rest of the brain.

### Neural timescales and sleep

To study how the hierarchical structure of neural timescales varies with brain state, we measured timescales during staged overnight sleep in 27 participants with a total of 2871 recording sites (see Table 1). Timescales fluctuated during sleep in correspondence with sleep stages (Figure 5a). We fit LME models with hierarchy and sleep stage and hierarchy-by-stage interaction and random intercepts for participant and recording site.

**Figure 5.**
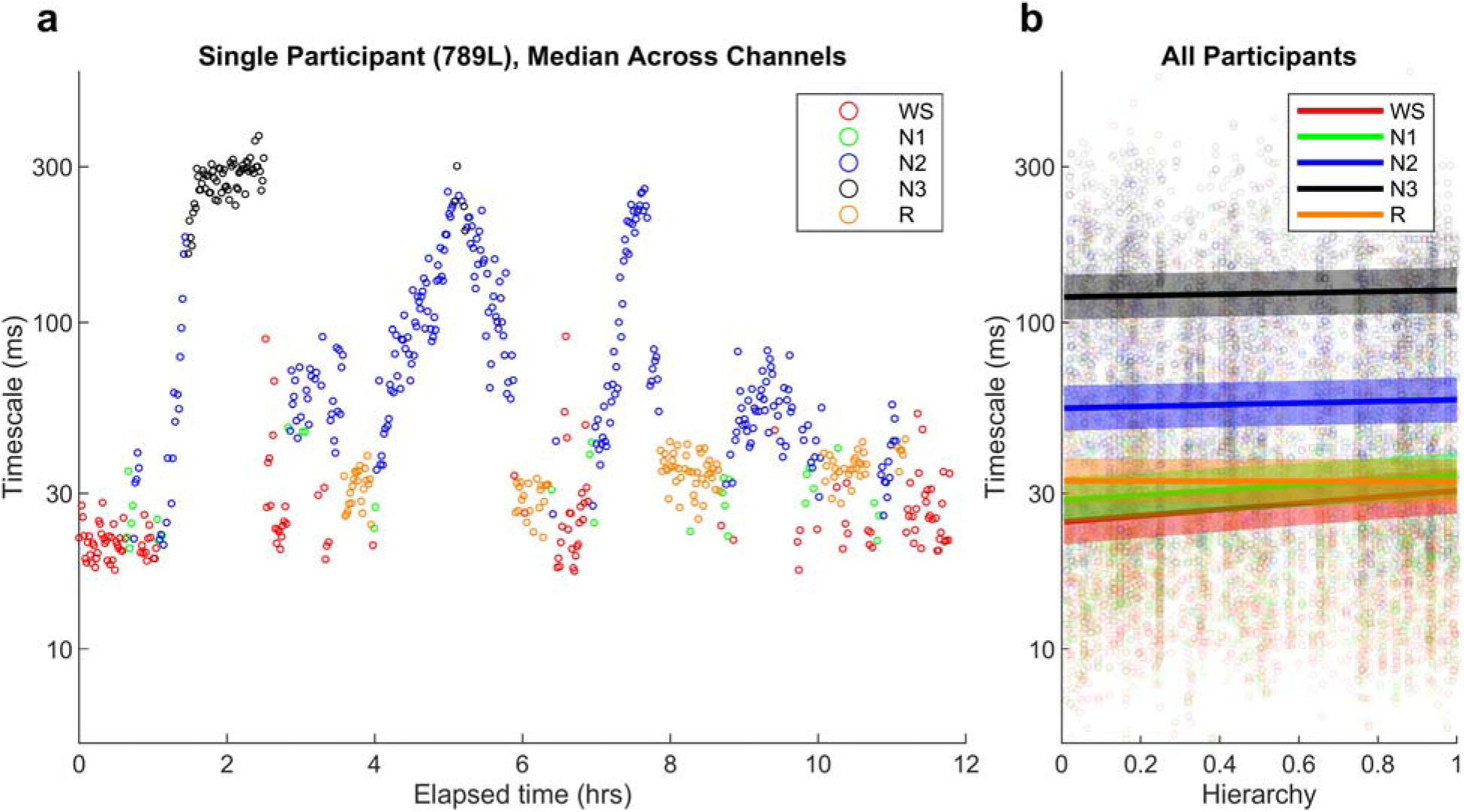
State-dependence of intrinsic timescales. **A.** Timescales vary with sleep stage. Data are from one night of sleep in an example participant. Each point represents the median timescale across sites during a one-minute segment of data. Intrinsic timescale is slower in N2 and N3 sleep compared to wake. **B.** Hierarchical organization of timescales changes with sleep stage. Each point is a single recording site in a color-coded sleep stage. Regression lines and 95% confidence intervals for the relationship with hierarchy in each stage are from a mixed effects model fit to data from all participants.

Timescales at Hierarchy = 0.5 (marginal predictions from LME model) were similar in magnitude during wakefulness (30 ms, 95% CI [25, 35]), N1 (34 ms, 95% CI [29, 40]), and REM sleep (36 ms, 95% CI [30, 42]), though the small differences were statistically significant from wake (N1, 1.1 fold, p < 0.001; REM, 1.2 fold, p < 0.001) due to large sample size at the recording site-level and within-unit structure of the comparisons (each recording site is observed in different sleep stages). Timescales slowed progressively in stages N2 (61 ms, 95% CI [52, 72]; 2.1 fold, p < 0.001 vs. wake) and N3 (134 ms, 95% CI [114, 156]; 4.5 fold, p < 0.001 vs. wake). This global slowing of timescales with NREM sleep depth parallels the overall reduction in network activity and the emergence of slow oscillatory dynamics during deeper NREM stages.

Sleep stages also affected the relationship between hierarchy and timescales (Figure 5b). During wakefulness and N1 sleep, a hierarchical gradient was present similar to wakeful resting-state recordings: timescales were faster in primary sensory and motor areas and slower in higher-order association areas (wake HTR = 1.25 [1.17, 1.33], N1 HTR = 1.19 [1.12, 1.27]; all as ratio [95% CI]; p = 0.17 wake vs. N1). In contrast, during N2, N3, and REM sleep, this gradient was reduced (N2 HTR = 1.06 [0.995, 1.13], N3 HTR = 1.05 [0.980, 1.13], REM HTR = 0.994 [0.929, 1.06]; all p < 0.001 relative to wake).

## Discussion

### Hierarchical organization of intrinsic timescales

A growing body of evidence establishes a hierarchical organization of intrinsic neural timescales across the cortex. The prevailing finding is that timescales are shorter in primary sensory and motor regions and longer in transmodal regions of the default mode, fronto-parietal, and cingulo-opercular networks (Murray et al., 2014). Evidence for a hierarchy of intrinsic timescales across the macaque cortex (Murray et al., 2014) used hierarchical position defined anatomically based on the laminar patterns of long-range corticocortical projections (Felleman and Van Essen, 1991). Complementary evidence comes from a large-scale computational model of the macaque cortex incorporating quantitative tract-tracing data and a gradient of excitatory synaptic strength across areas (Chaudhuri et al., 2015). The hierarchical organization of timescales was an emergent property in that model, arising from the combination of long-range connectivity architecture and local heterogeneity in recurrent synaptic strength, consistent with the results we show in Figure 4.

Several previous studies have characterized the hierarchical organization of neural timescales in human cortex. Some of those studies used an approach similar to that used here applied to resting-state BOLD fMRI data. For example, timescales were shown to lengthen systematically along the principal gradient, with slower timescales in transmodal regions (Ito et al., 2020). Similarly, topographic gradients of intrinsic dynamics were shown to mirror the principal gradient of functional organization (Shafiei et al., 2020), and ACW values measured in MEG data varied systematically between ‘core’ and ‘periphery’ regions defined using the first principal gradient (Golesorkhi et al., 2021a; Golesorkhi et al., 2021b; Wolff et al., 2022). By contrast, previous iEEG studies of this hierarchical organization relied on T1w/T2w ratio as a proxy measure (Gao et al., 2020), used established understandings of hierarchical relationships between cortical ROIs (Müller and Meisel, 2023), or used Euclidean anatomical distance (Cusinato et al., 2023).

Here, we introduce a formal method for determining hierarchical position of iEEG recording sites using DME (Figure 2). Unlike studies using BOLD fMRI data averaged across subject, the electrode-level resolution of iEEG allows the timescale gradient to be resolved within individual participants, as shown in the single-subject example in Figure 3. This approach builds directly on our prior work (Banks et al., 2023; Krause et al., 2023), in which we demonstrated that DME of iEEG resting-state functional connectivity reveals a geometrically interpretable, hierarchically organized embedding space. By using Human Connectome Project BOLD fMRI connectivity to generate the embedding and projecting iEEG recording sites into it via their respective MNI coordinates, we obtain hierarchical coordinates anchored to whole-brain functional organization.

### Network properties contributing to neural timescales

We show that neural timescales at the ROI level are strongly associated with two complementary measures of network topology, mean functional connectivity and a measure of network “hubness” (Figure 4). Together, these two panels suggest that the hierarchy of timescales reflects two partially separable properties of cortical network organization. Hubness captures the integrative role of an ROI within the network’s functional geometry, while mean connectivity reflects the overall influence that ROI has on network activity.

Our finding that timescales positively correlate with mean functional connectivity, i.e., ROIs with stronger average coupling exhibiting longer timescales, supports predictions of prior computational work based on macaque cortex (Chaudhuri et al., 2015; Gollo et al., 2015). In these models, stronger recurrent synaptic connections produced longer neural timescales, and regions with greater long-range connectivity (e.g., rich-club hubs) exhibited slower dynamics (Chaudhuri et al., 2015). Connectivity topology alone was thus sufficient to produce the timescale hierarchy.

The relationship between timescales and connectivity has been examined empirically in prior resting-state BOLD fMRI studies. A correlate of intrinsic neural timescales, lag-1 autocorrelation, was shown to positively correlate with mean FC across regions (Shinn et al., 2023). Longer timescales in resting-state BOLD fMRI have also been shown to associate with greater within-community functional connectivity, but the relationship with mean functional connectivity was weak (Lurie et al., 2024). Structural connectivity in humans and mice has been shown to positively correlate with τ_INT_ (Sethi et al., 2017; Fallon et al., 2020), consistent with the predictions of computational models (Chaudhuri et al., 2015; Gollo et al., 2015). Our Figure 4 (right panel) extends this line of evidence to functional connectivity in human iEEG, showing that regions with stronger mean functional coupling exhibit longer timescales. Together, these results indicate that the timescale gradient is related to both the structural scaffold and the functional connectivity patterns that arise from it.

Hubness, defined here as proximity to the center of embedding space, captures a different aspect of network architecture than connectivity magnitude alone. A region close to the embedding center has a connectivity profile to the rest of the brain that is maximally homogeneous (Banks et al., 2023), and thus well-suited to integrating information from a wide array of sources. In our previous work, we showed that regions such as anterior temporal cortex occupy hub-like positions in the iEEG embedding space. Here, we show that these regions also exhibit the longest timescales, consistent with the predictions of previous work (Gollo et al., 2015; Golesorkhi et al., 2021a). The correlation we observe between timescales and distance from the center of embedding space, our measure of hubness, provides a continuous, geometry-derived measure, enabling detection of graded variation across the cortical hierarchy.

### State-dependence of neural timescales

We show that timescales vary with sleep stage, i.e., fastest during wake, N1, and REM and progressively slower through N2 and N3, and that the hierarchical organization of timescales observed during wake and N1 is reduced during N2, N3, and REM, primarily due to an increase in timescale at the lowest levels of the hierarchy. Previous results for timescales derived from scalp EEG recordings showed a similar progression (wake<N1<N2≈REM<N3) (Zilio et al., 2021). Curiously timescales derived from the high gamma power signal in iEEG recordings were faster during sleep compared to wake (Müller and Meisel, 2023).

A previous study based on broadband iEEG signals and hierarchy defined by T1w/T2w ratio showed similar results to those presented here, with timescales increasing globally during NREM sleep and greatest changes in primary sensory areas (Cusinato et al., 2025). The authors attributed the latter observation to higher delta power in sensory areas during NREM sleep. Contrary to the results presented here, however, they found that the hierarchical organization of timescales persisted during NREM. The diversity of these results suggests that different methodological approaches, e.g., in the specific signal examined and the method for measuring hierarchy, may may capture distinct facets of state-dependent changes in cortical dynamics.

In the current study, as in most previous reports, our focus was on measuring the dominant timescale in each region. However, studies have shown that multiple timescales can be observed simultaneously in both field potentials and single unit data (Cavanagh et al., 2020; Lendner et al., 2024), and only some of the timescales are state-dependent (Lendner et al., 2024). This distinction may reflect additional structure in neural dynamics that captures variability within and between studies. We have previously shown, using DME of iEEG connectivity, that states of reduced consciousness are characterized by decreased functional differentiation and integration (Krause et al., 2023). The loss of timescale hierarchy in N2 and N3 observed here may similarly reflect loss of network differentiation, with the flattening of the timescale hierarchy representing the temporal-domain signature of the reduced network entropy.

REM sleep is electrophysiologically similar to wake, yet we find that timescale hierarchy is no longer detected during REM, similar to what we observe N2 and N3. This reinforces the view that aperiodic neural dynamics capture aspects of brain states that are not apparent in the overall power spectrum alone. The loss of hierarchy during REM may reflect the absence of task-relevant sensory input and attentional modulation, both of which have been shown to dynamically reshape timescale gradients in the awake brain (Gao et al., 2020; Zeraati et al., 2023; Raposo et al., 2025).

### Caveats and limitations

Several limitations should be considered when interpreting these results. As in all human intracranial recordings, participants had medically refractory epilepsy and may not be fully representative of a neurologically healthy population. We mitigated this concern by excluding recording sites identified as seizure foci from analysis, and we note that the timescale gradients found are consistent with that reported previously in non-patient cohorts. Because electrode placement was dictated solely by clinical considerations, coverage was sparse and non-uniform, no single participant sampled the full extent of the cortical hierarchy, and parts of cortex like the occipital lobe were systematically under-sampled. Anchoring recording sites to a group-level diffusion map embedding derived from Human Connectome Project fMRI partly offsets this limitation by providing a common hierarchical coordinate for sites pooled across participants. However, projecting individual electrode MNI coordinates onto a group, fMRI-derived embedding assumes a cross-modal spatial correspondence that is necessarily approximate.

Antiseizure medications are a further potential confound, as a higher drug load was shown to reduce temporal and spatial correlations in iEEG (Müller and Meisel, 2023). Because our participants were maintained on, and tapered from, these medications, drug effects could contribute to variance in timescales across participants and sessions. For comparisons across the hierarchy and between sleep stages, our repeated measurements within the same subject help cancel out variance added at the subject level from subject-level variations in medication, and we use a single recording session for resting-state or sleep, without making between-session comparisons.

In addition, our timescale estimates reflect a single dominant aperiodic timescale; recent work indicates that the aperiodic spectrum contains multiple coexisting timescales, only some of which are state-dependent (Lendner et al., 2024), a structure our approach does not resolve.

Our data include a mixture of subdural and sEEG recording sites, and we found slower timescales when measuring from sEEG versus subdural electrodes. We have previously reported post-surgical differences in delta power on subdural vs. sEEG recording sites (Dappen et al., 2025), although these differences mostly resolved by the recording period used in this study (about 1 week post-implant) and otherwise might have predicted the opposite result: that higher post-operative delta power in subdural electrodes might mean slower measured timescales. Though we account for hierarchy directly in our comparisons, there may be other confounding dimensions that cause electrode type differences through coverage differences.

Finally, sleep staging relied on a limited number of overnight scalp-EEG recordings, and seizures, medication, and the hospital environment may have influenced sleep architecture and arousal.

### Conclusions

In summary, we introduce a method for assigning hierarchical position to individual iEEG recording sites by projecting their MNI coordinates onto an axis derived from diffusion map embedding of resting state functional connectivity. We show that this measure of hierarchy captures the previously reported hierarchical organization of neural timescales, which we show are also organized by network connectivity and topology. We further show that the hierarchical organization is state-dependent, changing with sleep stage. This approach can be applied to any spatially resolved measure of neural activity, offering a framework for studying the organization of brain features across arousal and cognitive states, during development, and in clinical populations.

